# *Sf3b4* mutation in *Xenopus tropicalis* causes RNA splicing defects followed by massive gene dysregulation that disrupt cranial neural crest development

**DOI:** 10.1101/2024.01.31.578190

**Authors:** Casey Griffin, Kelsey Coppenrath, Doha Khan, Ziyan Lin, Marko Horb, Jean-Pierre Saint-Jeannet

## Abstract

Nager syndrome is a rare craniofacial and limb disorder characterized by midface retrusion, micrognathia, absent thumbs, and radial hypoplasia. This disorder results from haploinsufficiency of SF3B4 (splicing factor 3b, subunit 4) a component of the pre-mRNA spliceosomal machinery. The spliceosome is a complex of RNA and proteins that function together to remove introns and join exons from transcribed pre-mRNA. While the spliceosome is present and functions in all cells of the body, most spliceosomopathies – including Nager syndrome – are cell/tissue-specific in their pathology. In Nager syndrome patients, it is the neural crest (NC)-derived craniofacial skeletal structures that are primarily affected. To understand the pathomechanism underlying this condition, we generated a *Xenopus tropicalis sf3b4* mutant line using the CRISPR/Cas9 gene editing technology. Here we describe the *sf3b4* mutant phenotype at neurula, tail bud, and tadpole stages, and performed temporal RNA-sequencing analysis to characterize the splicing events and transcriptional changes underlying this phenotype. Our data show that while loss of one copy of *sf3b4* is largely inconsequential in *Xenopus tropicalis*, homozygous deletion of *sf3b4* causes major splicing defects and massive gene dysregulation, which disrupt cranial NC cell migration and survival, thereby pointing at an essential role of Sf3b4 in craniofacial development.

## Introduction

Nager syndrome (OMIM#154400) is a form of acrofacial dysostoses (AFD), a rare group of disorders characterized by malformations of the craniofacial skeleton and limbs (Halal et al., 1983). Patients with Nager syndrome specifically present with downslanting palpebral fissures, midface retrusion, micrognathia, defective middle ear ossicles, and hypoplastic or absent thumbs (Trainor and Andrews, 2013). Nager syndrome is a rare disorder, with approximately 100 reported cases. The craniofacial skeletal structures affected in Nager syndrome patients are neural crest (NC)-derived.

The NC is an embryonic cell-type unique to vertebrates. These cells arise from the neural plate border region of the developing embryos, undergo an epithelial-to-mesenchymal transition, migrate throughout the embryo, and eventually give rise to numerous cell-types and structures, including melanocytes, the peripheral nervous system in the trunk, and most of the craniofacial skeleton in the head. A subdomain of the NC, known as the cranial NC, originating from the mesencephalon and rhombencephalon, migrates as streams to populate the pharyngeal arches. The pharyngeal arches give rise to the majority of the differentiated tissues of the head and neck, with the core of NC in each arch developing into most of the craniofacial skeletal elements (Trainor and Krumlauf, 2000). The craniofacial structures affected in Nager syndrome patients are primarily derived from the first and second pharyngeal arches (Passos-Bueno et al., 2009).

The major cause of Nager syndrome, in approximately 60% of cases, is haploinsufficiency of the splicing factor 3b, subunit 4 (*SF3B4*) gene (Bernier et al., 2012; Czeschik et al., 2013; Petit et al., 2014). Rodriguez syndrome (OMIM#201170) is another condition due to mutations in *SF3B4* (Drivas et al., 2019). Although these patients share similar craniofacial abnormalities with Nager syndrome patients, the defects are usually more severe and involve lower limb anomalies (Rodriguez et al., 1990). *SF3B4* encodes SAP49, a component of the U2 subunit of the major spliceosome (Will and Luhrmann, 2011). The spliceosome is a complex made up of RNAs and proteins that functions to identify non-coding introns in precursor messenger-RNA (pre-mRNA) and promote accurate splicing at the surrounding splice sites. Recognition of the 5’ and 3’ splice-sites, as well as proper binding to the pre-mRNA and other parts of the spliceosome complex are required for the splicing process to occur. At each step of the splicing process, different sets of small nuclear ribonucleoproteins (snRNPs) are recruited, and the combination of RNA, small nuclear RNA (snRNA), snRNP, and non-snRNP protein interactions allow for splicing to occur at the proper location (Will and Luhrmann, 2011). SF3B4 contains two RNA recognition motifs at its N-terminal end through which it binds upstream to the branch site in the intronic region of the pre-mRNA, where it helps tether the U2 complex to the branch site.

Previous work in mouse models has shown that a loss of SF3B4 affects the axial skeleton and the forebrain (Yamada et al., 2020; Kumar et al., 2023), with a disruption in Hox gene expression and aberrant splicing of several chromatin remodelers (Kumar et al., 2023). In *Xenopus laevis*, morpholino-mediated knockdown of sf3b4 results in a loss of NC gene expression and causes a reduction or loss of craniofacial cartilages at tadpole stages through a mechanism that involves increased apoptosis (Devotta et al., 2016). A mutation in *sf3b4* in zebrafish has been reported to result in a phenotype reminiscent of retinitis pigmentosa, a spliceosomopathy affecting the photoreceptors in the retina (Ulhaq et al., 2023). Despite these efforts, for the most part the precise mechanisms underlying the pathogenesis of Nager syndrome remain largely unknown. It is especially puzzling that variants of a core component of this largely ubiquitous cellular machinery result in such an exquisitely cell-type and lineage-specific defect.

Here we report the characterization of a *Xenopus tropicalis sf3b4* mutant line generated using the CRISPR/Cas9 gene editing technology. We show that in the *sf3b4* mutants, NC induction is unaffected. However, at the tailbud stage cranial NC cell migration is disrupted, and these animals show increase apoptosis in the cranial region. While homozygous animals fail to survive beyond the tadpole stage, heterozygous animals are phenotypically largely indistinguishable from wildtype tadpoles. RNA-sequencing (RNA-seq) and gene ontology (GO) analyses of embryos at different developmental stages reveal an increase in aberrant splicing events, followed by a massive dysregulation of gene expression in homozygous animals. We propose that Sf3b4 is critical for cranial NC cell migration and survival, and that dysregulation of these processes is caused by aberrant splicing events and associated changes in gene expression upon *sf3b4* mutation.

## Results

### Developmental expression of sf3b4

We first analyzed the developmental expression of *sf3b4* in *Xenopus tropicalis* embryos using whole-mount *in situ* hybridization (WMISH). *sf3b4* transcripts are first detected at the end of gastrulation (NF stage 12.5) in the dorsal anterior ectoderm (Fig. 1A, B). At the early neurula stage (NF stage 14), the *sf3b4* expression domain encompasses the neural plate and the neural plate border (Fig. 1C). As neurulation proceeds (NF stage 17), *sf3b4* transcripts remain broadly enriched dorsally, including the prospective brain and spinal cord (Fig. 1E), and in a domain that overlaps with the *sox10* expression domain in the NC territory (Fig. 1F). As development continues, *sf3b4* is detected in the head region, including the developing brain, spinal cord, eyes, and migrating NC cells (Fig. 1 G-I). At tailbud stages (NF stages 25-31), *sf3b4* expression is largely confined to the brain, eyes, otic vesicles, pharyngeal arches, and the tailbud (Fig. 1K-M). Control sense probe at stage 14 (Fig. 1D) and stage 25 (Fig. 1J) did not show any signal. The expression pattern of *X. tropicalis sf3b4* is similar to that of *X. laevis sf3b4* (Devotta et al., 2016). The *sf3b4* expression domain over time encompasses regions of the embryo that are broader than the NC territory, therefore this expression pattern cannot account for the cell-type specific activity of sf3b4 in cranial NC and its derivatives.

**Figure 1:**
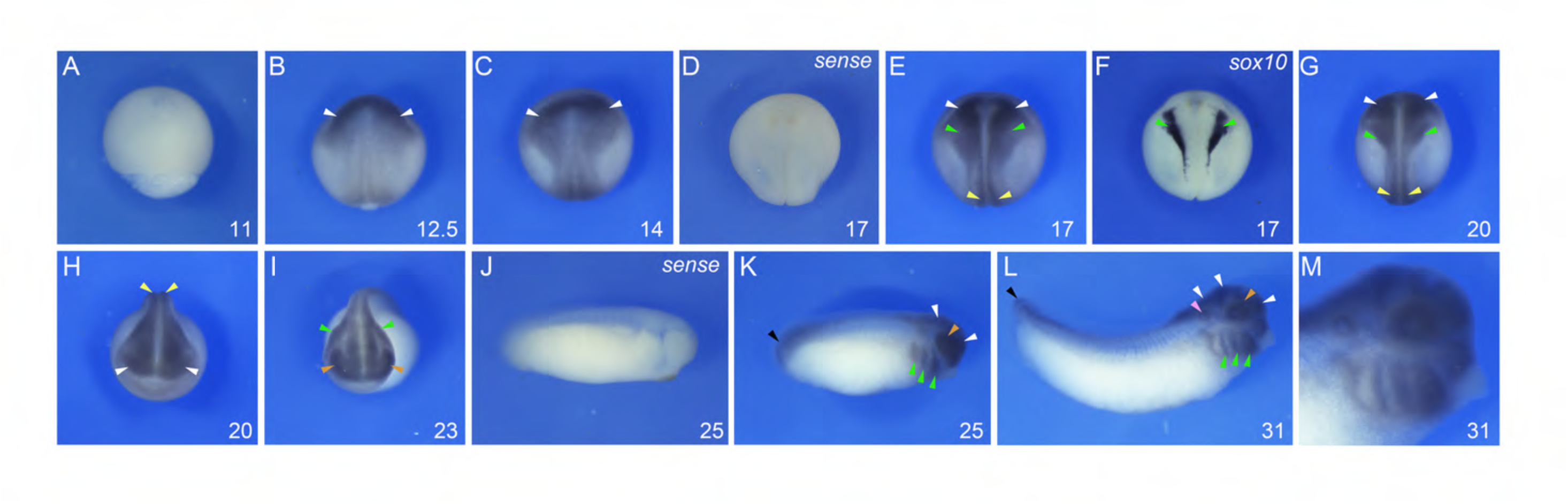
Developmental expression of *Xenopus tropicalis sf3b4.* *sf3b4* is not expressed at the mid-gastrula stage (**A**) but is first detected at the end of gastrulation in the dorsal ectoderm (**B**). During neurulation (**C, E-H**) *sf3b4* transcripts are detected in the developing neural plate/tube and neural crest forming regions, where it overlaps with *sox10* (**F**). At tailbud stages (**K-M**), *sf3b4* expression persists in the brain, eyes, migrating neural crest cells, and is also detected in the otic vesicles and the tailbud. Anterior neural plate/developing brain (white arrowheads), prospective spinal cord (yellow arrowheads), neural crest (green arrowheads), developing eyes (brown arrowheads), otic vesicle (purple arrowhead), and tailbud (black arrowheads). Embryos hybridized with a sense control are shown for stage 17 (**D**) and stage 25 (**J**). (**A-G**) Dorsal views, anterior to top. (**H-I**) Frontal views, dorsal to top. (**J-M**) Lateral views, anterior to right, dorsal to top. The embryonic stages (NF) are indicated in the lower right corner of each panel.

### Generation of a Xenopus tropicalis CRISPR/Cas9 sf3b4 mutant line

To generate this custom mutant line, five sgRNAs were synthesized targeting the first three exons of *X. tropicalis sf3b4* (Fig 2A). Mutations induced here disrupt both Sf3b4 RNA recognition motif domains. Of the embryos injected at the one cell stage, 20 reached adulthood. After reaching sexual maturity, one female was outcrossed to a wild type male and tested for germline transmission. Genotyping of the corresponding F1 embryos showed both -5 bp and -31 bp mutations. The remaining F1 embryos were reared and genotyped as adults through hindlimb web punch sampling. Out of 50 F1 adults, 15 had a confirmed -31 bp mutation. The -31 bp heterozygous F1 *sf3b4* adults were intercrossed to generate F2 mutants used for this study. The -31 bp mutation disrupts the start codon of Sf3b4, resulting in a frameshift at the first amino acid, and introducing an early stop codon 29 amino acids downstream (Fig 2B).

**Figure 2:**
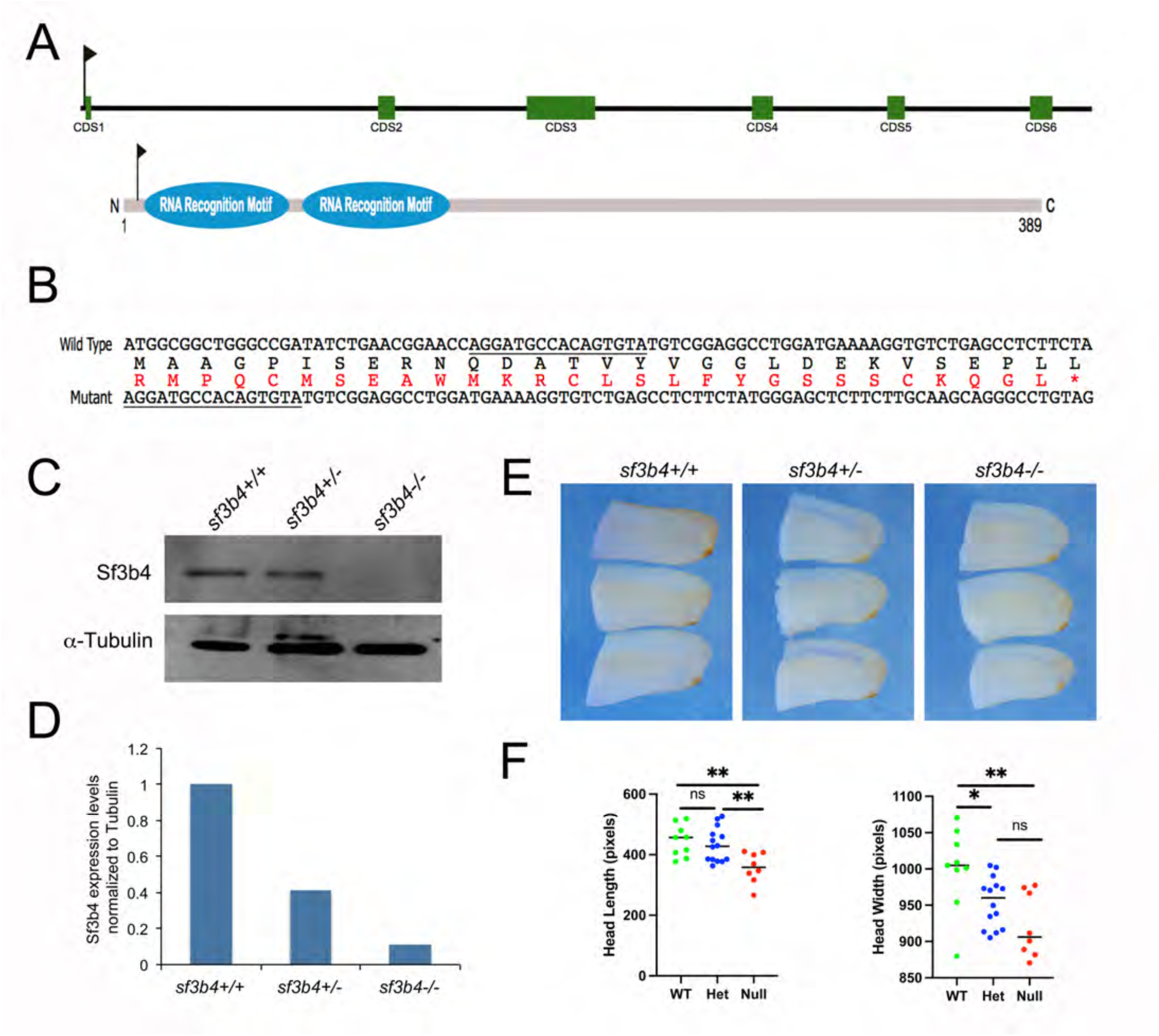
Generation of a *Xenopus tropicalis* CRISP/Cas9 *sf3b4* mutant line. (**A**) Schematic representation of the genomic *sf3b4* locus, and Sf3b4 protein. The relative position of the sgRNA target sequence is indicated (flags). CDS1-CDS6; Exon 1 to Exon 6. (**B**) Alignment of wildtype and mutant nucleotide and protein sequences. The underlined regions define where the wildtype and mutant sequences are identical after the 31bp deletion. A frameshift in the mutant nucleotide sequence results in an incorrect protein sequence (red) with an early stop (*) after amino acid 28. (**C**) Western blot analysis of protein extracts from wildtype (+/+), heterozygote (+/-), and homozygote (-/-) F2 tailbud stage embryos (NF stage 25), using an anti-Sf3b4 antibody. a-Tubulin is shown as a loading control. (**D**) ImageJ quantification of the Western blot. (**E**) Sf3b4 mutant embryos at stage 25 have reduced head length and width as compared to wildtype. Lateral views, anterior to right, dorsal to top. (**F**) Graph plotting the length and width for all three genotypes. ns = not significant, * = p<0.05, ** = p<0.01. Welch’s two-tailed unpaired t-test.

Western blot analysis of stage 25 wildtype (WT), heterozygous (Het), and homozygous (Null) mutant embryos shows that Sf3b4 expression levels are consistent with each genotype and the number of intact copies of the gene (Fig 2C, D). By gross morphology, mutant embryos at the tailbud stage (NF stage 25) display reduced head length and width, a phenotype that is especially more pronounced in Null mutants (Fig. 2E, F).

### Characterization of the Xenopus tropicalis CRISPR/Cas9 sf3b4 mutant embryos

Because NC-derived craniofacial skeletal elements are primarily affected in Nager syndrome patients, we analyzed by WMISH the expression of genes to visualize the NC at pre-migratory (*snai2, sox10, tfap2e*) and migratory (*sox9, sox10*) stages. Across all three genotypes, we found no difference in the expression of *snai2*, *sox10*, and *tfap2e* at early neurula stage (NF stage 14/15; Fig. 3A-B; Supp Table 1). Because *sf3b4* is also expressed in the neural plate (Fig. 1), we examined the expression of *sox2* at early neurula stage. This gene is unaffected in mutant embryos as compared to sibling controls (Fig. 3A; Supp Table 1). Upon closure of the neural plate (NF stage 20), *sox10* expression pattern in the NC territory remains unchanged across genotypes (Fig. 3C; Supp Table 1). At the migratory stage (NF stage 25) there is a notable phenotype in mutant embryos characterized by a decrease in the length of a subset of cranial NC streams, as evidenced by *sox9* and *sox10* expression (Fig. 4A; Supp Table 1). Quantification of the phenotype showed statistically significant reduction of the NC stream length in Het and Null embryos as compared to WT siblings (Fig. 4B). To understand the mechanism driving this phenotype, we performed TUNEL staining at different developmental stages. We found no change in apoptosis in *sf3b4* mutants at NF stage 20 (Supp Fig. 1), which correlates with the robust expression of *sox10* in the pre-migratory NC territory at this stage (Fig 3C). However, at the migratory stage (NF stage 25), there is a significant increase in apoptosis in the head region of Het and Null embryos, with a greater number of TUNEL positive cells in the Null as compared to the Het animals (Fig. 4C-D; Supp Table 1). These results suggest that while NC cell formation is not affected at the neural plate border in sf3b4 mutants, their migration in the pharyngeal arches is impaired.

**Figure 3:**
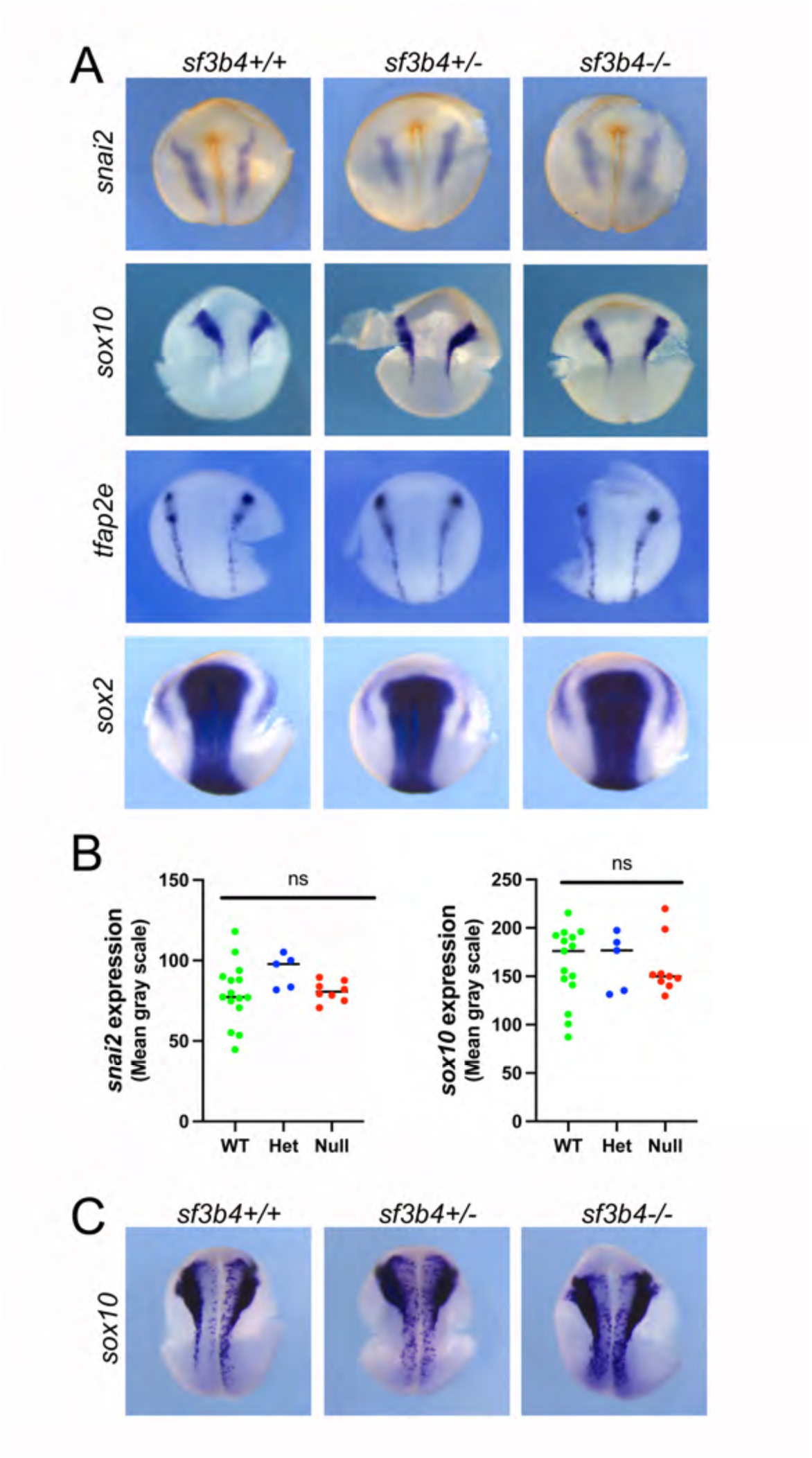
Molecular characterization of *sf3b4* mutant embryos at neurula stages. (**A**) At NF stage 14/15, the expression of *snai2*, *sox10*, and *tfap2e* in NC progenitors is largely unaltered in all three genotypes. The neural plate expression of *sox2* is also unaffected. sf3b4 wildtype (+/+), heterozygote (+/-), and homozygote (-/-). (**B**) ImageJ quantification of *snai2* and *sox10 WMISH signal*. ns = not significant. Welch’s two-tailed unpaired t-test. (**C**) At the end of neurulation, NF stage 20, the expression of *sox10* is largely unaltered in all three genotypes. (**A, C**) Dorsal views, anterior to top.

**Figure 4:**
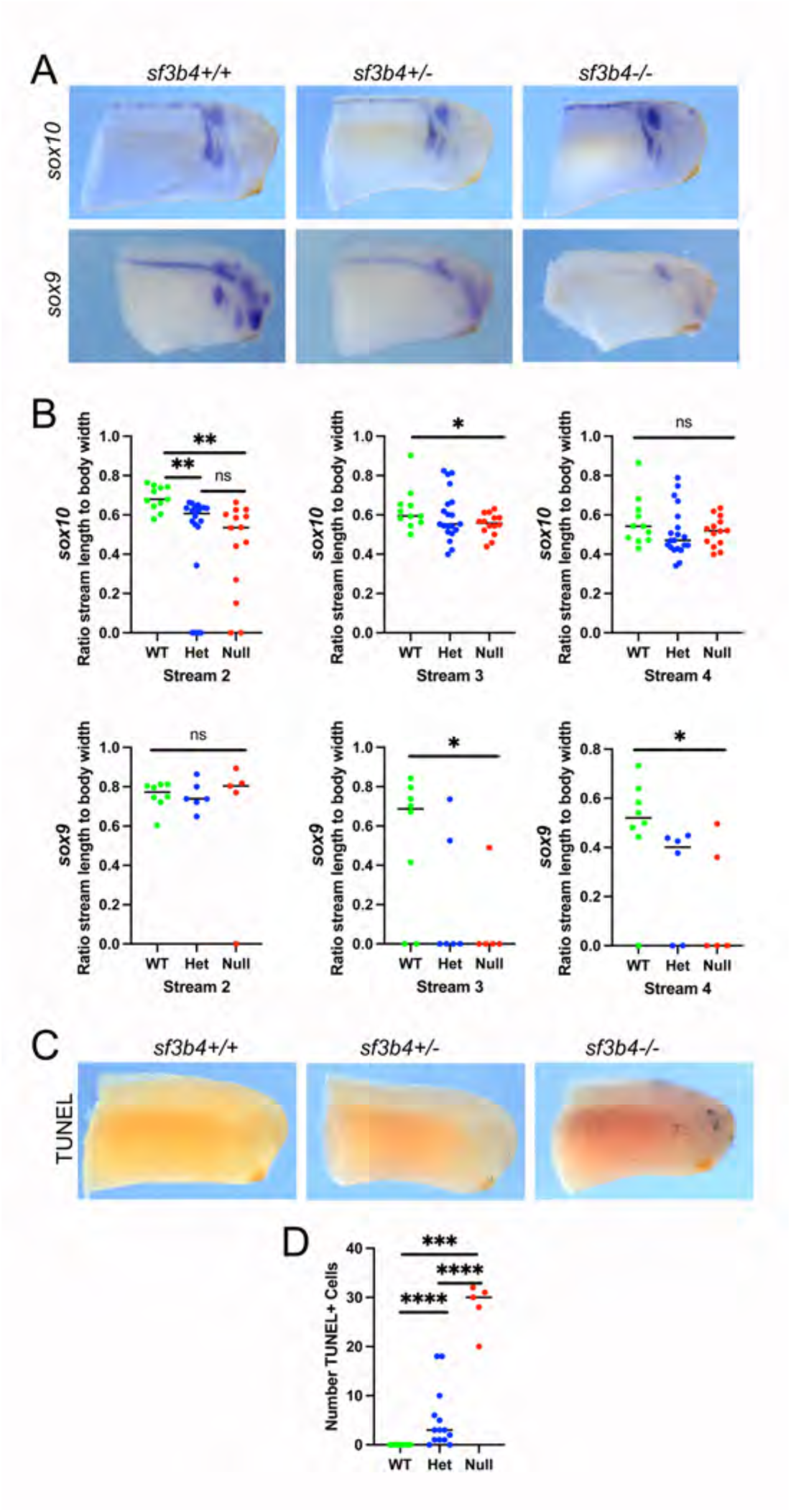
Molecular characterization of *sf3b4* mutant embryos at tailbud stage. (**A**) At NF stage 25, the expression of *sox9* and *sox10* in the NC streams is reduced or absent in heterozygous and null embryos. sf3b4 wildtype (+/+), heterozygote (+/-), and homozygote (-/-). (**B**) Quantification of stream length compared to full body width of embryos. ns = not significant, * = p<0.05, ** = p<0.01. (**C**) TUNEL staining of NF stage 25 embryos. (**D**) Quantification of TUNEL staining. *** = p<0.001, **** = p<0.0001. (**B, D**) Welch’s two-tailed unpaired t-test. (**A, D**) Lateral views, anterior to right, dorsal to top.

### Characterization of the Xenopus tropicalis CRISPR/Cas9 sf3b4 mutant tadpoles

We next analyzed mutant tadpoles at post-migratory stage (NF stage 40), when NC cells coalesce in the pharyngeal arches into foci of cartilage precursors expressing *sox9*. We noticed that the number of *sf3b4* Null animals recovered at this stage was lower than the expected mendelian ratio (Supp Fig. 2) suggesting that the phenotype described at the migratory stage may not be fully compatible with survival. Our analysis of the sf3b4 mutant tadpoles indicates that *sox9* expression is decreased in Null as compared to Het and WT embryos (Fig. 5A; Supp Table 1); however *sox9* expression is not completely lost in the mutants suggesting that the subset of NC cells that migrates into the pharyngeal arches can initiate a cartilage differentiation program. For comparison, Fig 5B shows a different batch of tadpoles stained for *sox9* expression with extended chromogenic reaction, presenting significant staining in the pharyngeal arches (Supp Table 1). We also noticed by gross morphology that all Null tadpoles appeared to present heart defects as compared to WT and Het animals (Fig. 5A,B), although this was not further investigated.

**Figure 5:**
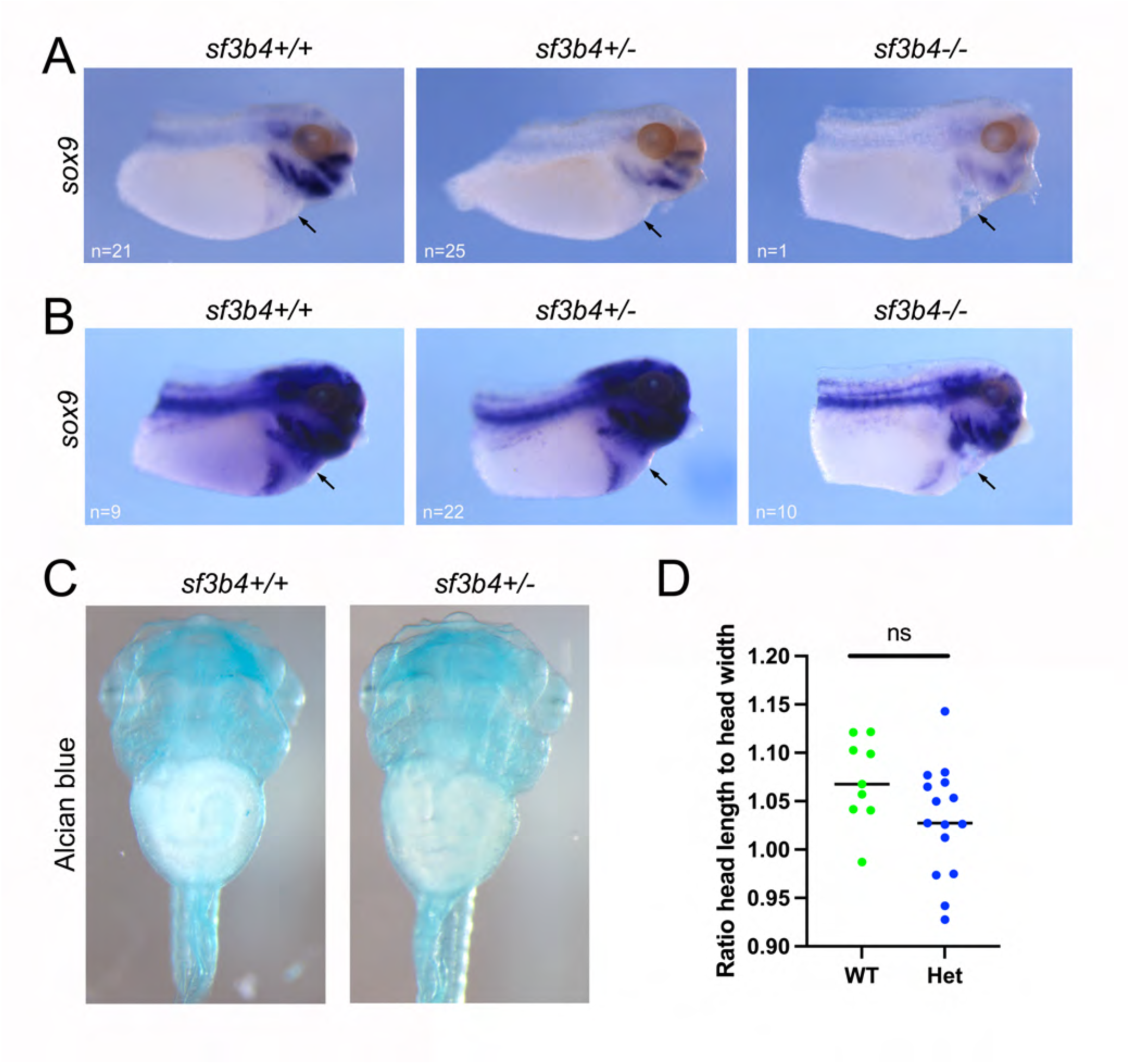
Molecular characterization of the *sf3b4* mutant tadpoles. (**A**) At NF stage 40, *sox9* expression in the branchial arches is reduced in heterozygote and null embryos. (**B**) Second set of tadpoles stained for *sox9* expression with extended chromogenic reaction, reveals robust overall sox9 expression in the pharyngeal arches. (**A, B**) Lateral views, anterior to right, dorsal to top. n, number of embryos analyzed per genotype. sf3b4 wildtype (+/+), heterozygote (+/-) and homozygote (-/-). (**C**) Alcian blue stained tadpoles, NF stage 45. Ventral views, anterior to top. sf3b4 wildtype (+/+) and heterozygote (+/-). (**D**) Graph plotting the ratio head length to head width for wildtype (+/+) and heterozygote (+/-) tadpoles. ns = not significant. Welch’s two-tailed unpaired t-test.

Finally, we analyzed the long-term consequences of *sf3b4* deletion on craniofacial cartilage formation at NF stage 45 by performing alcian blue staining. Interestingly, we did not recover any Null tadoples at this stage; all genotyped animals were either WT or Het (Supp Fig. 2), suggesting that the defects observed at earlier developmental stages are too severe for survival. Interestingly, head cartilage staining of WT and Het tadpoles reveals that these structures are largely unaffected in Het tadpoles, with a slight decrease in the overall size of the head, which is not statistically significant (Fig. 5C-D).

### Changes in gene expression associated with Sf3b4 loss of function

To identify gene networks disrupted in *sf3b4* mutants that may underly the Null phenotype observed, we performed bulk RNA-sequencing (RNA-seq) on pools of whole embryos from NF stages 15, 25, and 35 comparing WT, Het, and Null at each stage. Volcano plot analyses indicate that differential gene expression was very minimal at NF stage 15 across genotypes (Fig. 6A; top row). Furthermore, WT and Het samples analyses at all three stages show little to no differences (Fig. 6A; left column), suggesting that loss of one copy of *sf3b4* is largely inconsequential at these stages. By contrast pairwise comparison of Null vs. WT and Null vs. Het at NF stage 25 and stage 35 show a significant number of differentially expressed genes (Fig. 6A; middle and right columns). We next used Venn diagrams to determine the extent to which gene expression changes in Null vs. WT and Null vs. Het samples overlap at these 2 stages (Fig. 6B,C). Interestingly, the majority of differentially expressed genes in Null vs. Het largely overlap with that of Null vs. WT at stage 25 (Fig. 6B) and stage 35 (Fig. 6C), although the overall number of differentially expressed genes was much greater at stage 35 (2823 genes) than at stage 25 (352 genes) (Fig. 6C). Gene Ontology (GO) analyses for Biological Processes revealed an enrichment for terms such as “Wnt signaling pathway” and “mRNA splicing, via spliceosome” for downregulated genes, and terms like “cell division” and “DNA repair” for upregulated genes at NF stage 25 (Fig. 6D; Supp Table 2). Whereas at NF stage 35, “neural crest cell migration”, “negative regulation of extrinsic apoptotic signaling” and “mRNA splicing, via spliceosome” terms were enriched for downregulated genes, and “cell cycle”, “exocytosis” and “visual perception” terms enriched for upregulated genes (Fig. 6E; Supp Table 3). Taken together, these results demonstrate that gene expression changes in WT vs. Het are very minimal at all stages examined, suggesting a limited impact associated with the loss of one copy of *Sf3b4*, whereas when both genotypes are compared to their Null counterpart a similar dysregulation of gene expression is observed, and significantly more pronounced at stage 35.

**Figure 6:**
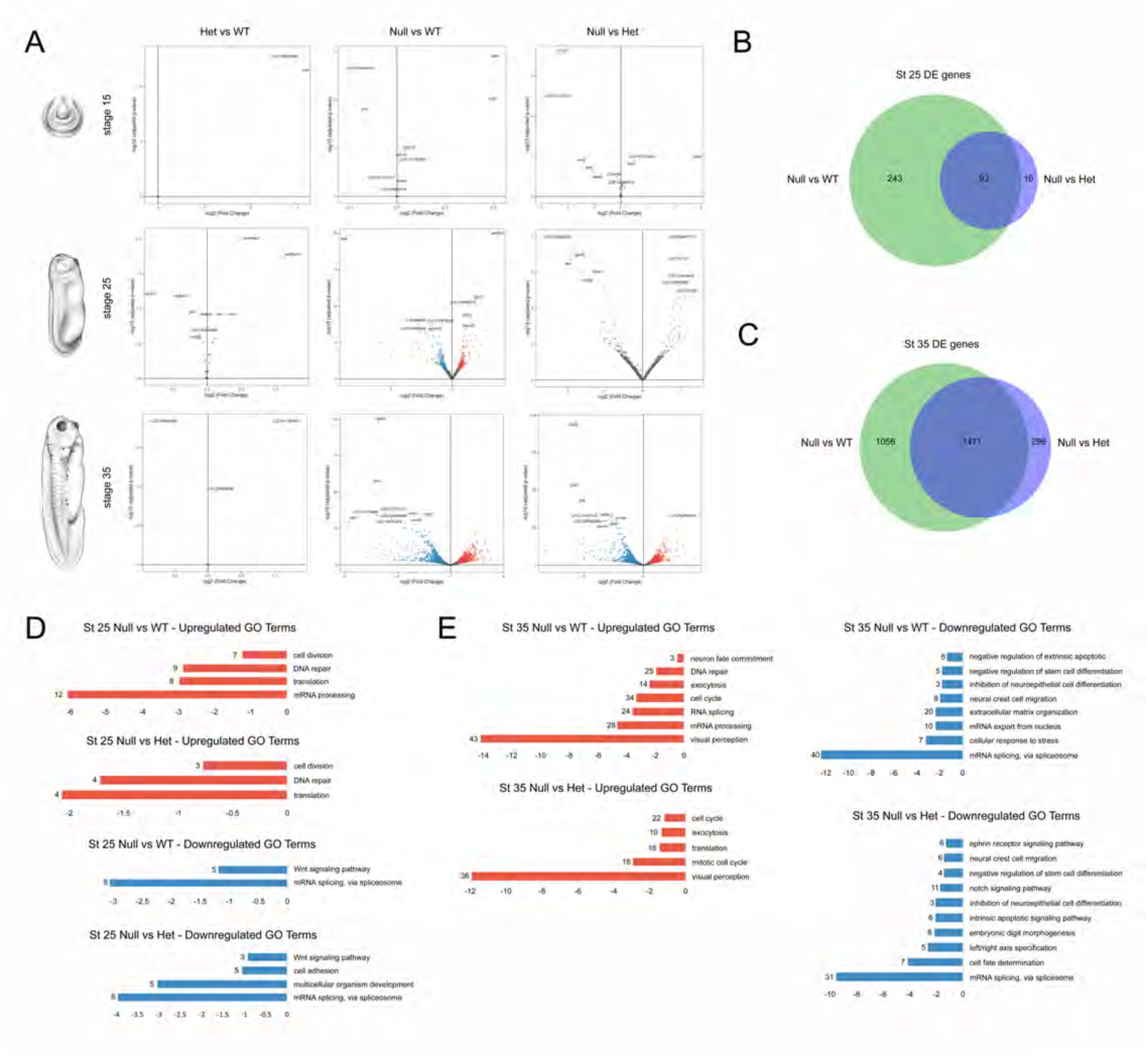
RNA-seq analysis of differentially expressed genes in wildtype and *sf3b4* mutant embryos. (**A**) Volcano plots showing significance (y-axis; log10 - adjusted p-value) vs. amplitude (x-axis; log2 - fold change) of differentially expressed genes across genotypes and stages. Genotype comparisons (Het vs. WT, Null vs. WT and Null vs. Het) are indicated on the top and embryonic stage (stage 15, 25 and 35) on the left. The top 10 genes for each comparison are labeled with their Xenbase ID. Xenopus illustrations @ Natalyan Zhan (2022). (**B-C**) Venn diagram analysis of differentially expressed genes at stage 25 (**B**) and stage 35 (**C**), comparing Null vs Wt and Null vs Het. Gene ontology (GO) term analysis of differentially expressed genes at NF stage 25 (**D**) and stage 35 (**E**) embryos. Upregulated terms are in red, downregulated terms in blue.

### Altered splicing events associated with Sf3b4 loss of function

Because Sf3b4 is an active component of the splicing machinery, we next examined the number and type of splicing events occurring at each stage (NF stage 15, 25 and 35), comparing each genotype, and focusing on four main events: skipped exons, retained introns, and 3’ and 5’ alternative splice sites (Fig. 7A). As observed for differentially expressed genes (Fig. 6A), we found very little difference between WT and Het when compared to Null at all three stages examined (Fig. 7B; Supp Fig 3). However, in all cases, there is a marked increase in the number genes with abnormal skipped exon events, especially at stage 25 (Fig. 7B; Supp Fig 3). Venn diagram analysis highlights the considerable overlap in genes with aberrant skipped exon between WT and Het when compared to Null at stage 25 (Fig. 7C) and stage 35 (Fig. 7D), with a greater overall number of genes affected at stage 25 (970 genes) than at stage 35 (180 genes) or stage 15 (140 genes) (Fig. 7C, D; Supp Fig. 3). GO analysis for Biological Processes at stage 25 shows an enrichment for terms such as “RNA splicing”, “DNA repair”, “regulation of embryonic development” and “regulation of apoptotic process” among others (Fig. 7E; Supp Table 4), while at stage 35, these terms include “mRNA splicing, via spliceosome”, “clathrin-dependent endocytosis”, “mRNA export from nucleus” and “ribosome assembly” (Fig. 7F; Supp Table 5). These results indicate that the complete loss of Sf3b4 function affects the spliceosome activity by preferentially promoting atypical skipped exon events more widely at stage 25 than at stage 35, while the loss of one copy of *sf3b4* had limited impact on spliceosome activity at both stages.

**Figure 7:**
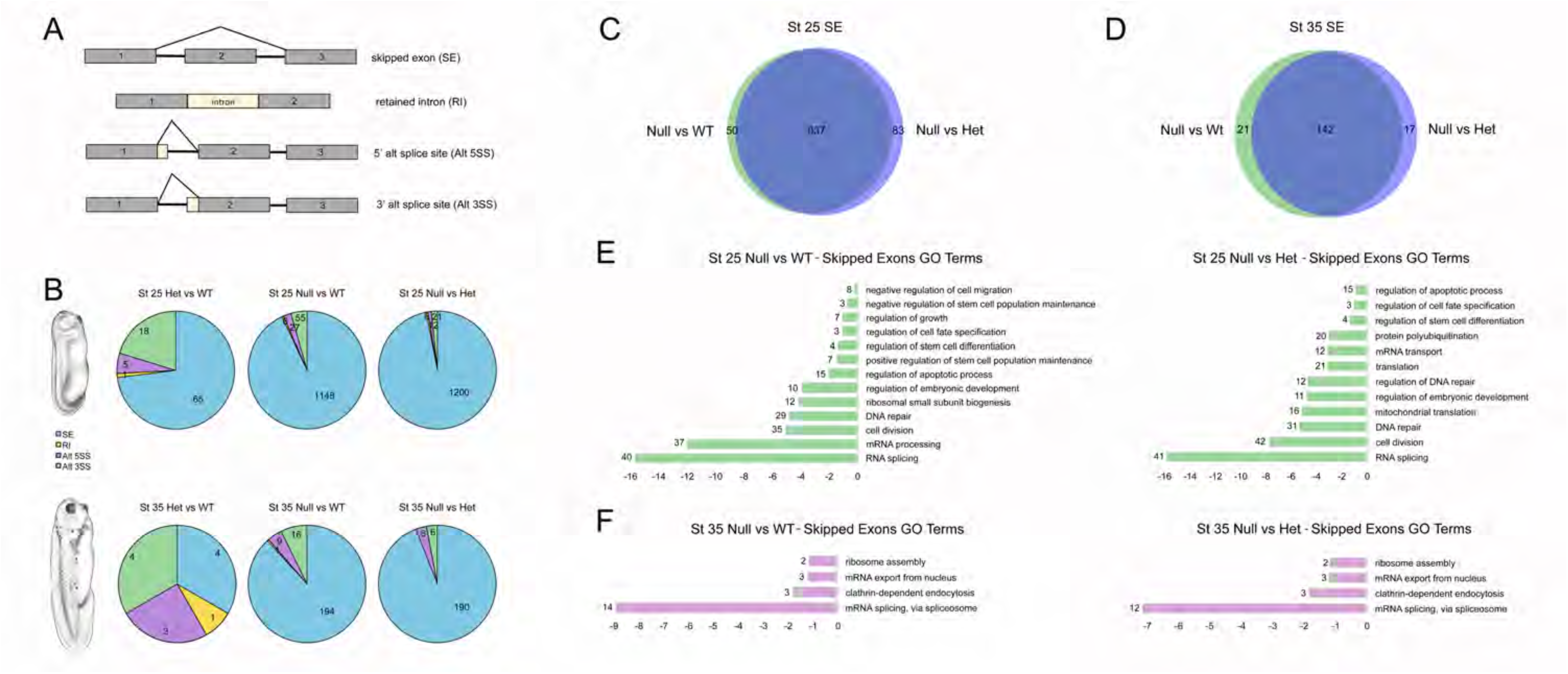
RNA-seq analysis of splicing events in wildtype and *sf3b4* mutant embryos. (**A**) Diagram of splicing events considered in this analysis. (**B**) Pie charts showing the type and number of splicing events occurring for each genotype comparison (Het vs. WT, Null vs. WT and Null vs. Het) at stage 25 and stage 35. Xenopus illustrations @ Natalyan Zhan (2022). (**C**) Venn diagram for genes with skipped exon at NF stage 25 (**C**) and stage 35 (**D**). WT and Het samples show substantial overlap at both stages. GO term analysis for genes with skipped exon at NF stage 25 (**E**) and stage 35 (**F**).

## Discussion

Here we report the generation and molecular characterization of a *Xenopus tropicalis* CRISPR/Cas9 *sf3b4* mutant line, as a novel model to investigate the pathogenesis of Nager syndrome, a condition that affects NC-derived craniofacial structures. We show that in the absence of Sf3b4 function NC induction is not affected. However, NC cell migration is disrupted at the tailbud stage, and coupled with an increase in apoptosis in the head region, a phenotype that is much more pronounced in Null than in Het animals. While *sf3b4* Null animals failed to survive beyond the tadpole stage, Het tadpoles were undistinguishable from their WT siblings, forming largely normal craniofacial cartilages. Temporal RNA-seq analysis confirmed at the transcriptome level the limited difference between WT and *sf3b4* Het embryos, and revealed aberrant pre-mRNA splicing events in stage 25 Null embryos, most notably characterized by an increase in the number of genes with atypical skipped exons, followed by a massive dysregulation of gene expression at stage 35. GO analysis of differentially expressed genes in *Sf3b4* Null revealed an enrichment in terms pertaining to mRNA splicing, apoptosis, cell cycle, and NC cell migration. Altogether, our data indicates that *sf3b4* haploinsufficiency is compatible with normal development in *X. tropicalis*, while homozygous deletion of *sf3b4* is detrimental to cranial NC cell migration and survival.

Splicing factor variants have been implicated in numerous diseases, referred to as spliceosomopathies. They form a singular group of diseases due to the nature of their phenotype affecting often a single cell/tissue-type, despite the predicted ubiquitous activity of the spliceosome in all cell types of the body (Griffin and Saint-Jeannet, 2020; Beauchamp et al. 2020). The tissues primarily affected in these pathologies include the retina (retinitis pigmentosa), spinal cord (amyotrophic lateral sclerosis), bone marrow (myelodysplastic syndromes), limb and the craniofacial skeleton (Griffin and Saint-Jeannet, 2020). Understanding the cell/tissue-specificity of these phenotypes has been quite challenging.

Mouse embryos with heterozygous Sf3b4 mutation show axial skeleton and forebrain defects, while Sf3b4 homozygous deletion is embryonic lethal (Yamada et al., 2020). RNA-seq analyses of Sf3b4 heterozygous mutants reveal a disruption in *Hox* gene expression and aberrant splicing of several chromatin remodelers known to regulate *Hox* genes (Kumar et al., 2023). In *X. laevis*, morpholino-mediated knockdown of sf3b4 results in a reduction of NC gene expression at neurula and tailbud stages, causing a reduction of NC-derived craniofacial cartilages at tadpole stages through a mechanism that involves apoptosis (Devotta et al., 2016). We did not observe a similar reduction in NC gene expression in the Sf3b4 *X. tropicalis* mutant embryos. Although *X. laevis* morphant phenotype was partially rescued by expression of human SF3B4 (Devotta et al. 2016), we cannot exclude that the morpholino may still also have off-target effects. Regardless, in both models cell death appears to be a common root cause in the presentation of the phenotype. While *X. tropicalis sf3b4* Het embryos exhibit a mild NC phenotype at tailbud stages, they appear to regulate/compensate this defect as they show virtually no craniofacial differences with WT animals at tadpole stage. Our data clearly indicate that loss of Sf3b4 in *X. tropicalis* manifests differently than in humans. The *sf3b4* Het embryos do not show an overall robust disease phenotype, and in many respects are much more similar to WT than Null embryos. This is further supported by our RNA-seq data (see below). Therefore, in this model of Nager syndrome, it is the Sf3b4 Null embryos that more closely recapitulate the human disease phenotype.

Our RNA-seq analyses at 3 different stages of NC development, pre-migratory (NF stage 15), migratory (NF stage 25) and post-migratory (NF stage 40), indicate that there is very little change in gene expression between WT and *sf3b4* Het animals (Fig. 6A), suggesting that loss of one copy of *sf3b4* is largely inconsequential in *X. tropicalis*, which is consistent with the phenotypes of these embryos (Fig. 5). When both genotypes are individually compared to *sf3b4* Null, we observed a similar pattern of gene dysregulation, with a much greater number of genes affected at stage 35 than at stage 25 or stage 15 (Fig. 6B, C; not shown). GO analyses of these gene sets revealed several terms linked to “mRNA splicing, via spliceosome”, “apoptosis” and “neural crest cell migration” enriched in genes downregulated at stage 35, while the most statistically significant GO term enriched among upregulated genes was “visual perception”. Genes downregulated related to “mRNA splicing, via spliceosome” include several genes that have been linked to other craniofacial spliceosomopathies, EFTUD2 (Lines et al., 2012), TXNL4 (Wieczorek et al., 2014) and PHF5A (Harms et al., 2023). We also found multiple genes causative of retinitis pigmentosa (a retina-specific spliceosomopathy, characterized by degeneration of photoreceptors), including PRPF3, PRPF8 and PRPF31 (Tanackovic et al. 2011), and SNRNP200 (Zhao et al., 2009). This is consistent with a recent study describing a photoreceptor phenotype in *sf3b4* mutant zebrafish larvae (Ulhaq et al., 2023). Here, we can speculate that the downregulation of the retinitis pigmentosa genes may indirectly cause the retinal phenotype, and the associated upregulation of genes linked to visual perception and retina function also reported in this study (Ulhaq et al., 2023). Also downregulated are genes related to “apoptosis”, including CDKN1 (cyclin dependent kinase inhibitor 1), a negative regulator of p53 (el-Deiry et al., 1994) and TP53 (tumor protein 53) itself (Chipuk et al., 2003). This is significant, as previous work has implicated apoptosis in craniofacial malformations underlying loss of Eftud2, Snrpb, and Txnl4a in *X. laevis* (Park et al., 2022), as well as the specific involvement of the p53 pathway in EFTUD2 mutant mice (Beauchamp et al., 2021), pointing at a common mechanism underlying craniofacial spliceosomopathies. Under the “neural crest cell migration” GO term, we recovered several members of the semaphorin family SEMA3B, SEMA4C, SEMA6A and SEMA6C, a semaphorin receptor (NRP1; neuropilin 1), EDNRB (endothelin receptor type B) and two transcription factors HIF-1a and TBX1. Most of these factors have been linked to NC development in several species, and pathogenetic variants of TBX1 and EDNRB cause two neurocristopathies, DiGeorge syndrome (Merscher et al., 2001) and Hirschsprung disease (Amiel et al., 1996), respectively. Future studies will determine whether the downregulation of these genes in *sf3b4* Null mutants underlies aspects of the NC-specific phenotype of this craniofacial spliceosomopathy.

The RNA-seq analyses combined with rMATS, a computational tool to detect splicing events, revealed very limited changes in the number and type of splicing events occurring between WT and *sf3b4* Het animals (Fig. 7B), again suggesting that one of copy of *sf3b4* in frogs is sufficient for spliceosome activity. When both genotypes are individually compared to *sf3b4* Null samples, a similar pattern of disrupted splicing events emerges, with a strong bias toward skipped exons, and a much greater number of genes affected at stage 25 than at stage 35 (Fig. 7C, D). This bias toward skipped exons could be due to the role of Sf3b4 in tethering the U2 complex to the branch site during splicing (Champion-Arnaud and Reed, 1994). In the absence of Sf3b4, the branch site stability may decrease, leading to the spliceosome falling off and exons skipping as alternate branch points are used instead. GO analyses identified several terms pertaining to “mRNA splicing”, “regulation of apoptotic process”, “DNA repair”, “cell division” and “regulation of embryonic development” enriched at stage 25, while at stage 35 the most statistically significant GO term enriched was “mRNA splicing, via spliceosome”, followed by “clathrin dependent endocytosis”. How these factors relate to NC and craniofacial development remain to be investigated.

The temporal transcriptomic analysis described here offers a unique perspective on the sequence of events and the mechanisms of Sf3b4 pathogenesis. Sf3b4 mutants show a massive disruption in spliceosome activity occurring primarily between NF stage 15 and stage 25, followed by a significant dysregulation of gene expression occurring around NF stage 35. This suggests that changes in gene expression are the likely consequence of spliceosome activity disruption and therefore may not be the primary cause of the phenotype, but rather the contributing factor. Future studies will focus on the plethora of genes mis-regulated in Null mutants at stage 35 (Supp Table 3) to identify the gene networks and pathways underlying this craniofacial condition.

## Materials and Methods

### Animal Care and Facility

The National *Xenopus* Resource (NXR; Woods Hole, MA) houses *X. tropicalis* as described (McNamara et al., 2018; Shaidani et al., 2020). Adult *sf3b4* -31/+ mutants were intercrossed via *in vitro* fertilization to produce embryos for this study. Females were given 20 U of Pregnant Mare Serum Gonadotropin (PMSG) (Bio Vender, Ashville, NC; Cat# RP17827210000) and 200 U of Human Chorionic Gonadotropin (hCG) (Bio Vender, Ashville, NC; Cat# RP17825010) to induce egg laying (Wlizla et al., 2018).

### Whole-mount in situ hybridization

Antisense digoxygenin-labeled probes (Genius kit; Roche, Indianapolis, IN) were synthesized using template cDNA encoding *X. tropicalis sf3b4* (Horizon; Cat# MXT1765-202789577) and *X. laevis sox10* (Aoki et al., 2003), *snail2* (Mayor et al., 1995), *tfap2e* (Hong et al., 2014), *twist1* (Hopwood et al., 1989) and *sox9* (Spokony et al., 2002). Whole mount in situ hybridization (WMISH) was performed as previously described (Harland, 1991).

### CRISPR/Cas9

Guide RNAs (sgRNAs) were designed utilizing CRISPRScan (https://www.crisprscan.org/) targeting the first three exons of *sf3b4* (Moreno-Mateos et al., 2015); T1: <u>GG</u>TGCCACGGTGTATGTCGG, T2: <u>GG</u>AACATGATAAAGCTCTAT, T3: <u>GG</u>GGTCTCTCATGATCTTGG, T4: <u>GG</u>CTTCGGACGCAGCCATTG, and T5: <u>GG</u>GGCGAGAAAATGGCGGCT. 5’ dinucleotides were converted to GG for increased mutagenic activity (Gagnon et al., 2014). SP6 MEGAscript kit (Ambion, Cat. No. AM1330) protocol was followed to synthesize all sgRNAs. Injections using the *X. tropicalis* Nigerian line (RRID: NXR_1018) at the one cell stage consisted of 500 pg of guide RNA and 1000 pg of cas9 protein into each embryo. One F0 female was outcrossed through IVF to a wild type male, resulting in F1 embryos with -5 bp or -31 bp mutations. The -31/+ (*Xtr.sf3b4^emNXR^*, RRID: NXR_3056) *sf3b4* mutant line is available through the NXR (https://www.mbl.edu/xenopus).

### Embryo collection and genotyping of tissue samples

To genotype adult frogs, tissue samples were taken from webbing on hindlimbs using single-use biopsy punches (VWR 21909-140). Heterozygous mating pairs were intercrossed to produce embryos for *in situ* hybridization and RNA-seq analyses. Embryos collected for *in situ* hybridization were fixed in MEMFA (10 mL 10X MEMFA salts, 10 mL 37% formaldehyde, 80 mL NF H_2_O) overnight at 4°C. Embryos were stored long term at -20°C in 100% ethanol. Tail clips for genomic DNA (gDNA) extraction were taken from fixed embryos after rehydration in PBS. Tissues to be used in RNA-seq analysis were immediately preserved on dry ice and stored long term at -80°C.

gDNA extractions were done using Qiagen DNeasy Blood & Tissue Kit (Qiagen, Valencia, CA; Cat# 69506). PCR amplification was done using the following primers for targeted region: forward primer 5’-AATGAAACACCCTCTATGCGC-3’ and reverse primer 5’-AGAGATGGAGCCTGCACC-3’. PCR product was then purified by following NucleoSpin PCR Clean-up procedure (Macherey-Nagel; Cat# 740609.250) and sent for Sanger sequencing to confirm genotypes.

### Western blot analysis

Pools of 10 embryos were homogenized in lysis buffer and concentrated, and western blot analysis was performed as previously described (Devotta et al., 2016). Primary antibodies were as follows: anti Sf3b4 polyclonal antibody (Proteintech; Cat# 10482-1-AP; 1:2000 dilution) and anti a-tubulin antibody (Sigma Aldrich; Cat# T9026; 1:500 dilution). Secondary antibodies were anti-rabbit (EMD Millipore; Cat# MAB201P) and anti-mouse IgG (Abcam, Cat# ab6820) coupled to horseradish peroxidase (1:10,000 dilution).

### Quantification of in situ hybridization signal

The *in situ* hybridization signal was measured using ImageJ software. The image was reverted to an 8-bit gray image and inverted. The area of staining was selected, and the mean gray scale was calculated for each area. The left and right sides of each embryo were calculated separately and then averaged together for the final value. Two-tailed Welch’s unpaired t-test was performed using Prism Graphpad (v10.01) to determine statistical significance.

### Quantification of migratory streams

The length of neural crest streams was measured using ImageJ software. Each stream length was measured and the ratio of stream length to total embryo length was calculated. Two-tailed Welch’s unpaired t-test was performed using Prism Graphpad (v10.0.1) to determine statistical significance.

### TUNEL assay and quantification

TUNEL staining was carried out as previously described (Hensey and Gautier, 1998; Devotta et al., 2016). To quantify cell death, embryos were imaged individually, and a set region of the head was chosen in which the number of TUNEL-positive cells were manually counted for each embryo. Two-tailed Welch’s unpaired t-test was performed using Prism Graphpad (v10.0.1) to determine statistical significance.

### Cartilage staining

Tadpoles at NF stage 45 were fixed in MEMFA overnight at room temperature. Embryos were then moved to Alcian Blue solution (60 mg Alcian Blue, 30 mL acetic acid, 70 mL 100% ethanol) for 2.5 days. Embryos were rinsed in 1% HCl in 70% ethanol for 2 days. After rehydration, embryos were bleached in 1% H_2_O_2_, 5% Formamide in 1X SSC buffer and imaged.

### RNA isolation and sequencing

Pools of 5 embryos were used for RNA extraction via the Qiagen RNeasy Micro Kit (Qiagen, Valencia CA; Cat# 74004). RNA was eluted in RNase-free water and sent to the New York University Genome Technology Center for RNA-sequencing. RNA QC/QA was performed on a bioanalyzer before performing automated stranded RNA-seq library preparation with polyA selection. Samples were run on the Illumina NovaSeq 6000 system with SP 100 cycle flow cells.

### RNA-Sequencing data analysis

RNA-seq data were analyzed by sns rna-star pipeline (https:igordot.github.io/sns/routes/rna-star.html). Adapters and low-quality bases were trimmed using Trimmomatic (v0.36) (Bolger et al. 2014). Sequencing reads were mapped to the reference genome (XENTR_10.0, https://download.xenbase.org/xenbase/Genomics/JGI/Xentr10.0) using the STAR aligner (v2.7.3) (Dobin et al., 2013). Alignments were guided by a Gene Transfer Format (GTF) file, which was converted from a General Feature Format (GFF3) file. The mean read insert sizes and their standard deviations were calculated using Picard tools (v.2.18.20) (http://broadinstitute.github.io/picard). The genes-samples counts matric was generated using featureCounts (v1.6.3) (Liao et al., 2014), normalized based on their library size factors using DEseq2 (v1.30.1) (Love et al., 2014), and differential expression analysis was performed. The Read Per Million (RPM) normalized BigWig files were generated using deeptTools (v.3.1.0) (Ramirez et al., 2016). To compare the level of similarity among the samples and their replicates, we used two methods: principal-component analysis and Euclidean distance-based sample clustering. All the downstream statistical analyses and generating plots were performed in R environment (v4.0.3) (https://www.r-project.org/).

rMATS (v4.0.2) (Shen et al., 2014) was used for detecting differential alternative splicing events from RNA-seq data. Events with a mean of inclusion junction counts (IJC) or a mean of skipped junction counts less than 10 were removed. Only events with an inclusion level difference of more than 0.1 or less than -0.1, with a p-value < 0.05 were included.

The results of GO analysis were generated by DAVID 2021 (Huang et al., 2007) using gene lists from differential expressed genes and splicing events. Venn diagrams were generated using DeepVenn (Hulsen, 2022).

## Acknowledgements

We thank members of the Saint-Jeannet laboratory past and present for their support and helpful discussions, and Benjamin McKenzie for technical help. We are grateful to the Genome Technology Center (GTC) for expert library preparation and RNA sequencing. The work benefited from the support of Xenbase (http://www.xenbase.org/ - RRID:SCR_003280).

## Funding

This work was supported by grants from the National Institutes of Health to J-P. S-J. (R01-DE025468), C. G. (F32-DE030699) and to M. H. (P40-OD010997 and R24-OD030008).

## Competing Interests

The authors declare no competing interests.

## Data Availability Statement

All data supporting the results are available from the corresponding author upon request. The RNA-seq data have been deposited onto GEO database under accession number GSE249075.

## Notes

### Competing Interest Statement

The authors have declared no competing interest.

