## Supplemental Data for "*Sf3b4* mutation in *Xenopus tropicalis* causes RNA splicing defects followed by massive gene dysregulation that disrupt cranial neural crest development"

**Supplementary Materials**

**Supp Fig. 1: TUNEL staining at neurula stage (NF Stage 19/20) comparing the 3 genotypes.**

(**A**) Neurula stage embryo at NF stage 19/20. Xenopus illustrations @ Natalyan Zhan (2022). (**B**) TUNEL staining of NF stage 20 embryos shows no difference across genotypes. In all panels, dorsal view, anterior to top.

**Supp Fig. 2: Genotype frequency at different stages of development.**

**Supp Fig. 3: RNA-seq analysis of splicing events in wildtype and *sf3b4* mutant embryos at stage 15. (A)** Pie charts showing the type and number of splicing events occurring for each genotype comparison (Het vs. WT, Null vs. WT and Null vs. Het) at NF stage 15. Xenopus illustrations @ Natalyan Zhan (2022). (**B**) Venn diagram for genes with skipped exon at stage 15 (**C**). WT and Het samples show substantial overlap. (**C, D**) GO term analysis for genes with skipped exon at stage 15.

**Supp Table 1: Developmental gene expression analysis per genotypes.**

**Supp Table 2: GO analysis of differentially expressed genes in *sf3b4* mutants at stage 25**

**Supp Table 3: GO analysis of differentially expressed genes in *sf3b4* mutants at stage 35**

**Supp Table 4: GO analysis of genes with skipped exons in *sf3b4* mutants at stage 25**

**Supp Table 5: GO analysis of genes with skipped exons in *sf3b4* mutants at stage 35**

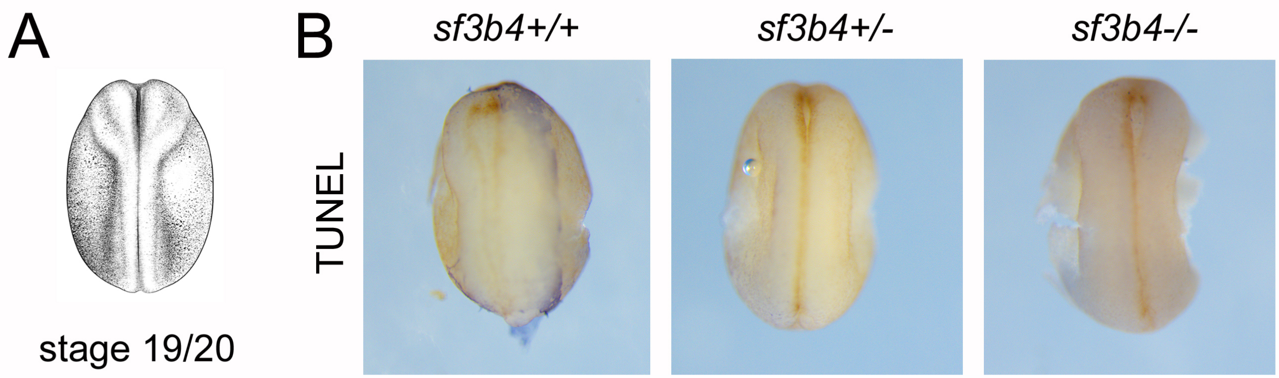


Supp Figure 1

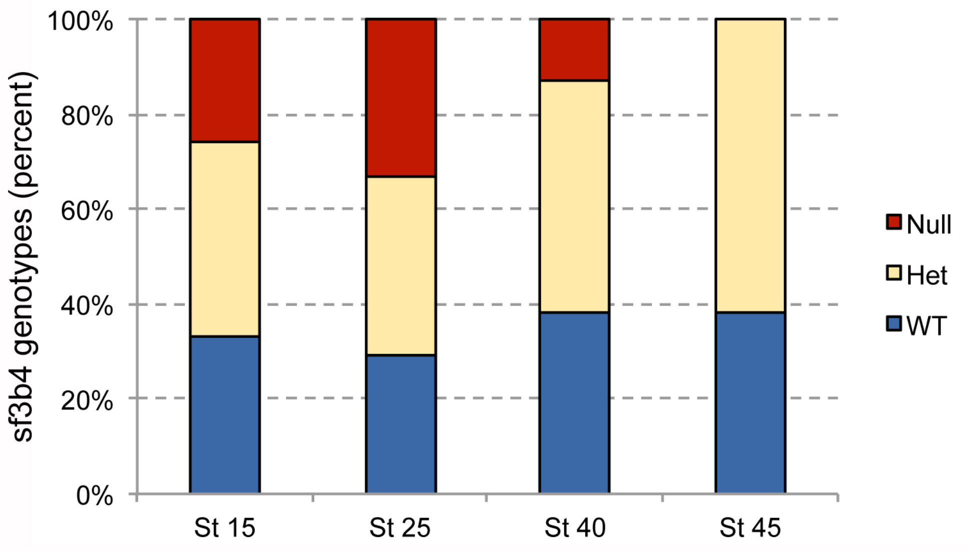


Supp Figure 2

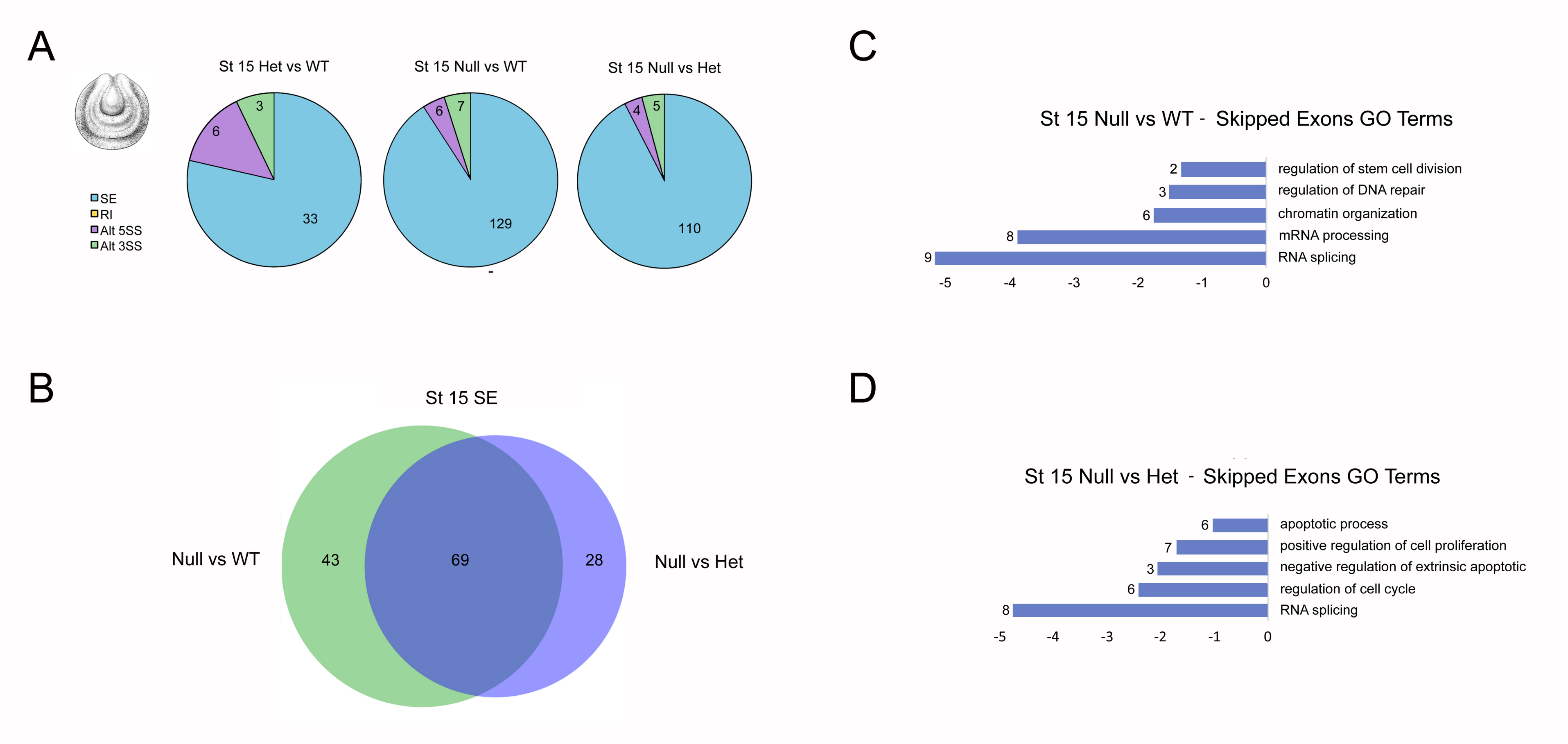


Supp Figure 3

| ISH Probe | Stage (NF) | WT | Het | Null |
|---|---|---|---|---|
| snai2 | 15 | 11 | 10 | 6 |
| | 15 | 5 | 5 | 3 |
| sox10 | 15 | 5 | 9 | 5 |
| | 15 | 10 | 11 | 6 |
| | 15 | 15 | 13 | 9 |
| tfap2e | 15 | 3 | 11 | 7 |
| sox2 | 15 | 5 | 8 | 6 |
| sox10 | 20 | 3 | 8 | 4 |
| sox9 | 25 | 8 | 8 | 8 |
| sox10 | 25 | 2 | 6 | 6 |
| | 25 | 9 | 14 | 8 |
| Sox9 | 40 | 10 | 16 | 0 |
| | 40 | 21 | 25 | 1 |
| | 40 | 10 | 8 | 7 |
| | 40 | 9 | 22 | 10 |

**Supp Table 1: Developmental gene expression analysis per genotypes**
